# Community member, community liaison and researcher perspectives on the creation of HapMap and 1000 Genomes Project iPSCs

**DOI:** 10.64898/2026.01.09.698717

**Authors:** Neda Gharani, Peter Ikhane, Tatyana Pozner, Sonia Sampson, Matthew W. Mitchell, Clement Adebamowo, Laura B. Scheinfeldt

## Abstract

The National Human Genome Research Institute (NHGRI) Repository for Human Genetic Research (NHGRI Repository) distributes renewable lymphoblastoid cell lines (LCL) and cell line DNAs donated by members of 26 communities who have taken part in the International HapMap and 1000 Genomes Projects. The relatively recent introduction of induced pluripotent stem cell (iPSC) reprogramming technologies offers a new opportunity to create iPSCs from HapMap and 1000 Genomes Project biospecimens. However, since relevant iPSC technology did not exist when the HapMap and 1000 Genomes Projects began, it was not explicitly included in the consent process. To explore donor community and researcher perspectives on potential HapMap and 1000 Genomes Project iPSC creation, we conducted an online, semi-structured survey. The majority of community (86%) and research (86%) respondents agreed that creating iPSCs from HapMap and 1000 Genomes Project biospecimens could benefit the research community; however non-trivial concerns (36% community respondents and 16% researcher respondents) and doubts (21% community respondents and 11% researcher respondents) about HapMap and 1000 Genomes Project iPSC creation were also shared. Given these results, we will continue to explore opportunities to engage with HapMap and 1000 Genomes Project communities that include educational materials and detailed discussions regarding the potential for creating iPSCs from their community biospecimens. In parallel, we will continue to grow the existing NHGRI Repository iPSC collection with new donor submissions through the Human Pangenome Reference Consortium (HPRC) that explicitly consented to the creation of iPSCs.

## Background

The National Human Genome Research Institute (NHGRI) Repository for Human Genetic Research (NHGRI Repository) is a biorepository that stores and distributes renewable lymphoblastoid cell lines (LCL) and cell line DNAs as well as associated information donated by members of 26 communities living around the world who have taken part in the International HapMap Project ^1-6^ and the 1000 Genomes Project ^7-11^. These participants have given consent for their samples to be shared for a broad range of future research and for their genomic information to be publicly shared; however, the NHGRI Repository does not include any personal or medical information.

The NHGRI Repository, therefore, includes some of the most extensively characterized and diverse human cell and DNA resources available for research ^12,13^; notably, public whole genome sequencing data are available for the majority of the collection (n>3,200) ^10^. Only qualified researchers can access NHGRI Repository samples, they must submit a statement of research intent, and the researcher and their respective organization must sign a formal agreement to use the samples only for their specified research. Moreover, quarterly reports of NHGRI Repository sample usage are sent to community liaisons, including researchers, community members, and individuals that fulfill both roles, to distribute to the HapMap and 1000 Genomes Project communities that they have worked with. Each report documents researchers who have requested and have been given samples, the institution where they work, their statement of research intent as well as a lay summary of the intended research. Community liaisons are additionally asked to distribute a selection of scientific articles that have used HapMap and 1000 Genomes Project samples and data to further highlight how the collection is being used over time to support research on genetic and genomic variation, health and disease.

The introduction of induced pluripotent stem cell (iPSC) reprogramming technology ^14,15^ and subsequent expansion to include methodologies relevant to NHGRI Repository biospecimens ^16^ offer a new opportunity to reprogram human iPSCs from peripheral blood mononuclear cells (PBMCs) from HapMap and 1000 Genomes Project participants. This technology allows researchers to differentiate iPSCs into any cell type, including neurons, cardiomyocytes, and hepatocytes, which is an exciting new opportunity for research into human health and disease ^14,15,17-19^. Relevant iPSC technology ^16^ did not exist when the HapMap and 1000 Genomes Projects began, so community participants were not told that their samples could be used in this particular manner. While the original consent did allow researchers to use cell lines for many different types of research and as an unlimited source of DNA, creating different kinds of cells and tissues from their samples may not have been what participants expected ^12^. The potential for HapMap and 1000 Genome Project iPSC creation therefore raises ethical questions that involve donor autonomy, community engagement, potential for re-identification, and research use and misuse. This potential also raises broader questions about how best to honor the intentions and interests of culturally, geographically, and politically diverse communities whose donated samples now support global science, and how to dialogue with them and their representatives to ensure sustained trust and fairness in the research collaboration ^2,12^.

The DISCUSS Project has studied the potential to derive iPSCs from previously collected biospecimens and has outlined key areas for consideration, including the adequacy of the consent, participant expectations, the feasibility of recontact, and governance of downstream applications ^20^. Many patients and members of the public generally support iPSC research, but they have also expressed concerns about privacy, immortalization, cellular transformation, and the creation of gametes or embryos ^21,22^. More recently in 2021, NHGRI organized a meeting entitled *Ethical Issues Associated with the Creation and Use of Induced Pluripotent Stem Cell (iPSC) Lines Derived from NHGRI-Supported Sample Collections* that included iPSC experts, genomics experts, and ethical, legal and social implications (ELSI) experts, as well as researchers that have worked with HapMap and 1000 Genomes Project communities. The associated report (https://www.genome.gov/sites/default/files/media/files/2023-09/iPSC_Meeting_Report_Sept2023.pdf) acknowledges two key components needed for consideration of iPSC establishment that were highlighted by the DISCUSS Project: the NHGRI Repository governance process and the language for broad research activities included in the HapMap and 1000 Genomes Project consent documentation. In addition, the meeting included discussion of the potential for variation in societal and cultural concerns regarding iPSC usage. To better understand the ethical concerns regarding the potential creation of HapMap and 1000 Genomes Project iPSCs, we conducted an online semi-structured survey to collect feedback from the communities that participated in the HapMap and 1000 Genomes Projects as well as from researchers that have used the collection in their studies.

## Methods

We developed a semi-structured online survey to evaluate the perspectives of community members, community liaisons and researchers on the potential for creating HapMap and 1000 Genomes Project iPSCs. We used the SurveyHero web-based survey software to implement the survey. Survey email invitations were sent between January and July of 2025. All survey responses were private and anonymous; no location or IP information was collected from survey participants.

Emails were sent to each set of community liaisons, and each of these emails included a SurveyHero link for them to share with the community members they have worked with (**Supplementary File A** includes the survey given to the community liaisons and community members). Instructions on how to translate the survey content with Google Translate were included in each community survey invitation email. After the initial email invitation, two email reminders were sent at least one month apart. A separate SurveyHero link was emailed to all researchers that have received biospecimens from the NHGRI Repository (**Supplementary File B** includes the survey given to researchers). After the initial email invitation, two email reminders were sent at least one month apart.

Both questionnaires included four main structured questions related to the possibility of creating iPSCs from HapMap and 1000 Genomes Project samples as well as free text field options for sharing additional feedback. This survey study was approved by the Coriell Institutional Review Board (#193).

## Results

We received completed survey results from 14 community representatives (community liaisons, and community members) and other respondents that received a survey invitation from a community liaison (**Table 1**). Since we do not have direct access to community members but rather indirect access through community liaisons, the survey invitations were sent to community liaisons to share with community members. We therefore do not know the number of community members that were invited by community liaisons to participate in the survey and were not able to calculate a community member response rate. In addition, of the 14 community survey responses, we cannot confirm the extent to which these were from liaisons, participants, community members or other community representatives.

**Table 1.** Community Respondent Details.

| Community Descriptor | Abbreviation | Associated Project(s) | N in collection | N liaison respondents | N community member respondents | N liaisons and community member respondents | N other respondents | N total respondents |
| --- | --- | --- | --- | --- | --- | --- | --- | --- |
| African Ancestry in Southwest, USA | ASW | HapMap, 1000GP | 107 |  |  |  |  |  |
| African Caribbean in Barbados | ACB | 1000GP | 120 |  |  |  |  |  |
| Bengali in Bangladesh | BEB | 1000GP | 144 |  |  | 1 |  | 1 |
| British from England and Scotland | GBR | 1000GP | 100 |  |  |  |  |  |
| Chinese Dai in Xishuangbanna | CDX | 1000GP | 102 |  |  |  |  |  |
| Chinese in Metropolitan Denver, CO, USA | CHD | HapMap | 129 |  |  |  |  |  |
| Colombian in Medellín, Colombia | CLM | 1000GP | 136 |  |  |  |  |  |
| Esan in Nigeria | ESN | 1000GP | 173 |  |  |  |  |  |
| Finnish in Finland | FIN | 1000GP | 103 | 1 | 1 |  |  | 2 |
| Gambian in Western Division, Mandinka | GWD | 1000GP | 179 |  |  |  |  |  |
| Gujarati Indians in Houston, TX, USA | GIH | HapMap, 1000GP | 117 | 1 |  |  |  | 1 |
| Han Chinese in Beijing, China | CHB | HapMap, 1000GP | 162 |  |  |  |  |  |
| Han Chinese South, China | CHS | 1000GP | 163 |  |  |  |  |  |
| Iberian Populations in Spain | IBS | 1000GP | 157 | 1 |  | 1 | 6 | 8 |
| Indian Telugu in the UK | ITU | 1000GP | 118 |  |  |  |  |  |
| Japanese in Tokyo, Japan | JPT | HapMap, 1000GP | 131 |  |  |  |  |  |
| Kinh in Ho Chi Minh City, Vietnam | KHV | 1000GP | 124 |  |  |  |  |  |
| Luhya in Webuye, Kenya | LWK | HapMap, 1000GP | 122 |  |  |  |  |  |
| Maasai in Kinyawa, Kenya | MKK | HapMap | 205 |  |  |  |  |  |
| Mende in Sierra Leone | MSL | 1000GP | 128 |  |  |  |  |  |
| Mexican Ancestry in Los Angeles, CA, USA | MXL | HapMap, 1000GP | 104 |  |  |  |  |  |
| Peruvian in Lima, Peru | PEL | 1000GP | 122 |  |  |  |  |  |
| Puerto Rican in Puerto Rico | PUR | 1000GP | 139 |  |  |  |  |  |
| Punjabi in Lahore, Pakistan | PJL | 1000GP | 158 |  |  |  |  |  |
| Sri Lankan Tamil in the UK | STU | 1000GP | 128 |  |  |  |  |  |
| Toscani in Italia | TSI | HapMap, 1000GP | 117 |  |  |  |  |  |
| Yoruba in Ibadan, Nigeria | YRI | HapMap, 1000GP | 229 |  |  | 1 | 1 | 2 |

We received completed survey results from 125 researchers who have previously used NHGRI Repository biospecimens. We initially sent survey invitations to 2,673 researcher emails in our database, and the research response rate was 5%. Responses to the four main questions related to the possibility of creating iPCSs from HapMap and 1000 Genomes Project samples are detailed below.

### “Do you support the creation of iPSCs from HapMap and 1000 Genomes Project samples for research purposes”

As displayed in **Figure 1**, the majority of community (79%; 11/14 respondents to this question) and researcher (78%; 97/125 respondents to this question) survey responses to the question “Do you support the creation of iPSCs from HapMap and 1000 Genomes Project samples for research purposes” were “yes”. None of the community respondents and five (4%) researchers answered “no”.

**Figure 1.**
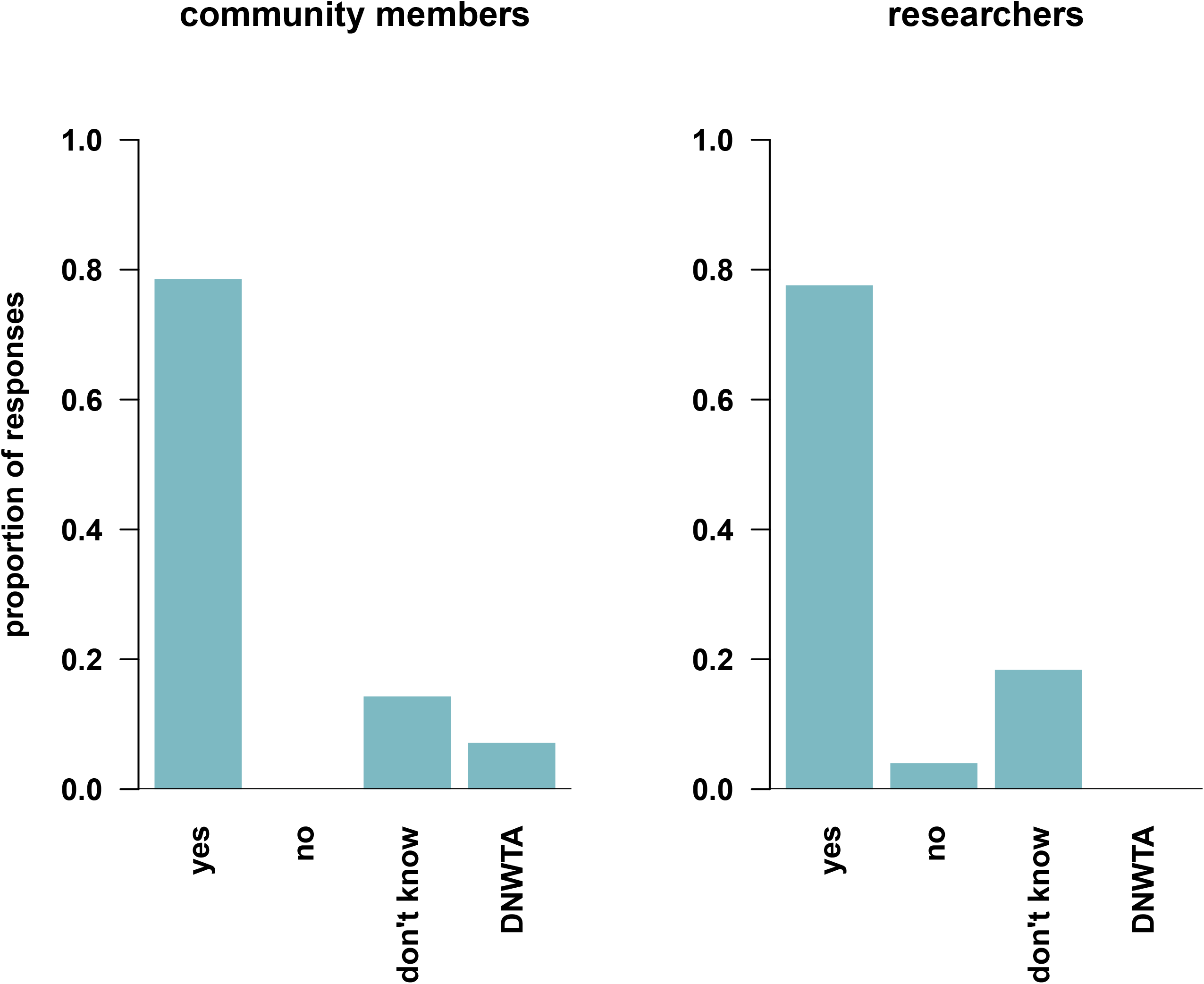
HapMap and 1000 Genomes Project iPSC Creation Support. Survey participants were asked the structured question “Do you support the creation of iPSCs from HapMap and 1000 Genomes Project samples for research purposes” with the following options: “yes”, “no”, “don’t know”, “do not want to answer”.

In addition to this multiple-choice question, all respondents were offered a free text field to share additional feedback to the following prompt: “Please let us know why you do or do not support the creation of iPSCs from HapMap and 1000 Genomes Project samples for research.” Several community liaisons and members shared feedback about the potential scientific benefits of iPSCs, lending their support to the creation of HapMap and 1000 Genomes Project iPSCs, while one community respondent, though supportive of iPSC creation, did express the need for proper safeguards and a second community respondent expressed concerns given the lack of explicit consent (see **Supplemental File C** for all free text answers).

Many researcher respondents shared enthusiasm for HapMap and 1000 Genomes Project iPSCs to benefit research given their diverse participants, the availability of genetic, genomic and transcriptomic characterizations, potential to characterize tissue-specific models to explore the functional implication of genetic variation, potential to reduce the need for animal models, and for use as reference materials (see **Supplemental File C** for more detail). Researcher concerns included the lack of consent for the specific use of biospecimens in the creation of iPSCs. One researcher pointed out that there are newer consented cell collections with explicit consent for iPSC creation, and another researcher expressed reservations about the creation of iPSCs without explicit consent, particularly when involving indigenous communities (see **Supplemental File C** for more detail).

### “Do you feel that there are potential benefits to adding iPSCs to the collection”

Figure 2. shows that the majority of community (86%; 12/14 respondents to this question) and researcher (86%; 107/125 respondents to this question) responses were “yes” to the question “ Do you feel that there are potential benefits to adding iPSCs to the collection”. Two (14%) community respondents and 16 (13%) researchers answered “Don’t know”, and two (<2%) researchers answered “no”.

**Figure 2.**
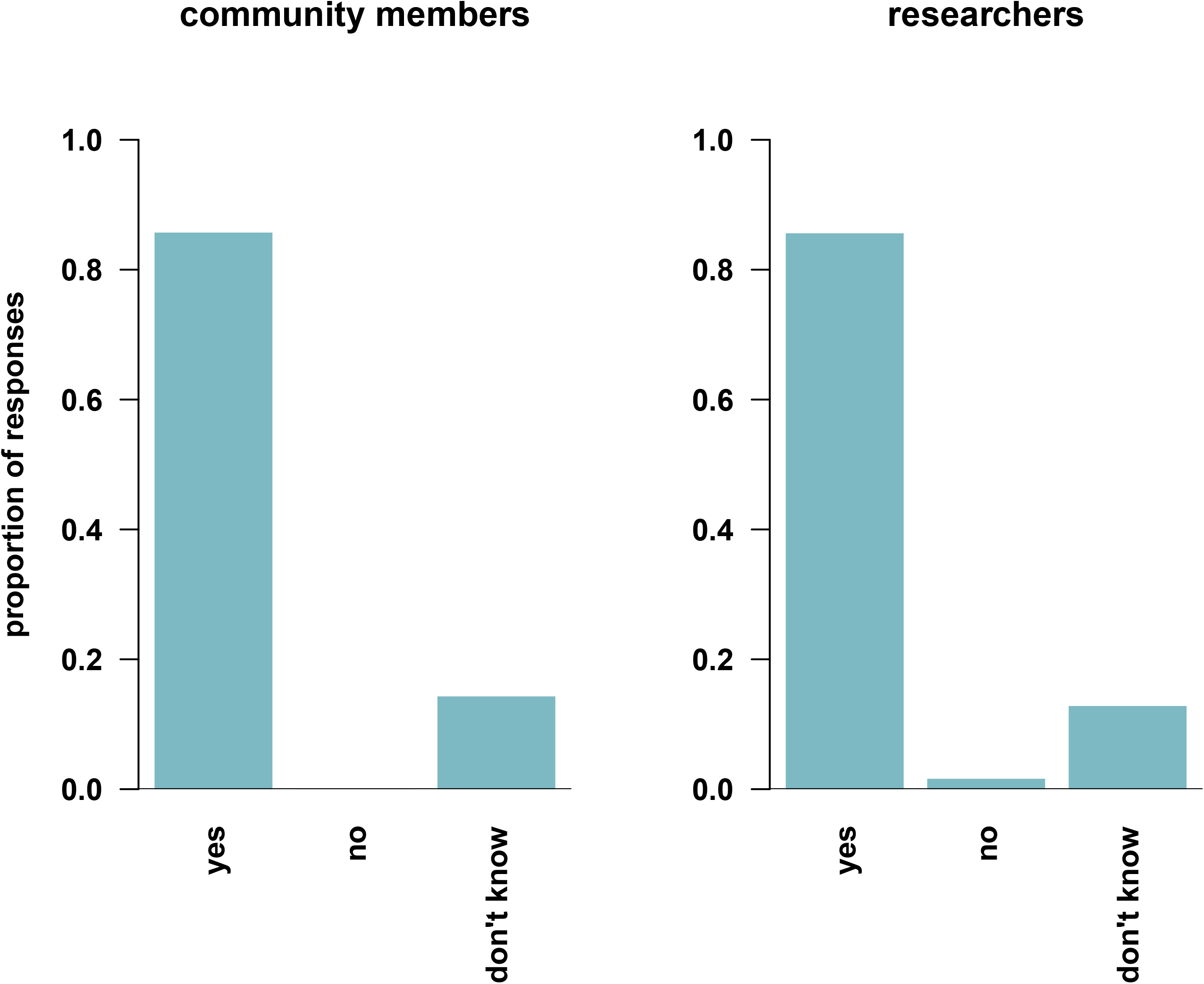
Potential iPSC Benefits. Survey participants were asked the structured question “Do you feel that there are potential benefits to adding iPSCs to the collection” with the following options: “yes”, “no”, “don’t know”, “do not want to answer”.

In addition to this multiple-choice question, all respondents were offered a free text field to share additional feedback to the following prompt: “Please share anything more about potential benefits to adding iPSCs into the collection.” Again, several survey responses added additional feedback involving the potential scientific benefit of HapMap and 1000 Genomes Project iPSCs (see **Supplemental File C** for more detail).

### “Do you have any concerns about adding iPSCs into the collection”

Figure 3. displays answers to the question “Do you have any concerns about adding iPSCs into the collection”. Five (36%) community respondents answered “yes”, six (43%) answered “no”, and three (21%) answered “don’t know”; 20 (16%) researcher respondents answered “yes”, 90 (72%) answered “no”, 14 (11%) answered “don’t know”, and one (1%) answered “do not want to answer”.

**Figure 3.**
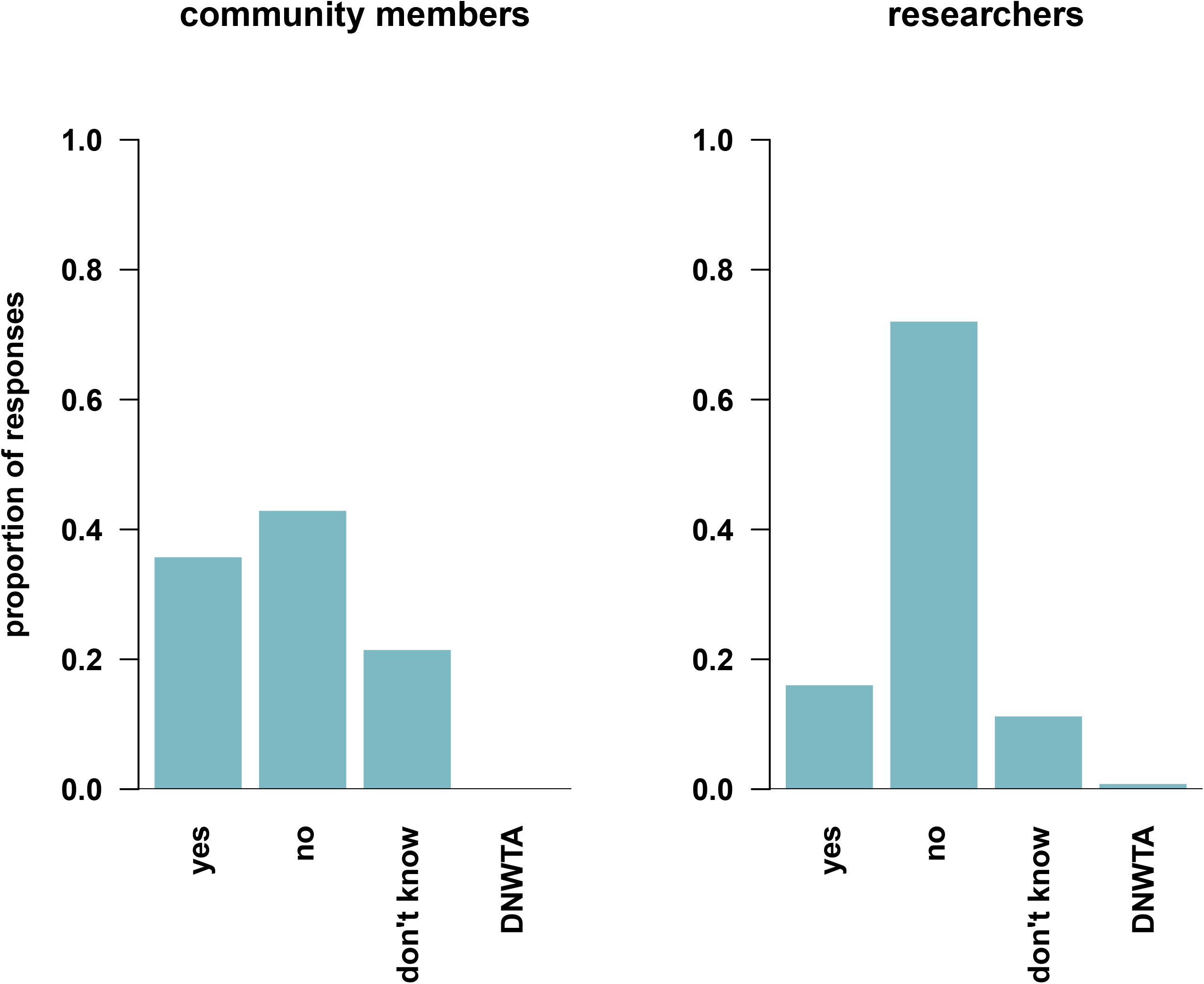
iPSC Concerns. Survey participants were asked the structured question “ Do you have any concerns about adding iPSCs into the collection “ with the following options: “yes”, “no”, “don’t know”, “do not want to answer”.

In addition to this multiple-choice question, all respondents were offered a free text field to share additional feedback to the following prompt: “Please share anything more about concerns with adding iPSCs into the collection.” Additional community comments focused on the lack of consent to create iPSCs (see **Supplemental File C** for more detail). The majority of additional researcher comments included the lack of consent to create iPSCs, that the original consent was for generating public genetic data but not phenotypic data and that iPSCs would generate phenotypic data, should iPSCs be created strict research safeguarding to prevent questionable uses is needed e.g. gamete production and possibility of allogeneic transplant, as well as concerns about exploitation of vulnerable communities, social discrimination, and public mistrust (see **Supplemental File C** for more detail).

### “Do you feel that you need more information about iPSCs before you can make an informed decision”

Figure 4. displays answers to the question “Do you feel that you need more information about iPSCs before you can make an informed decision”. Seven (50%) community respondents answered “yes”, five (36%) answered “no”, and two (14%) answered “don’t know”; 26 (21%) researcher respondents answered “yes”, 84 (67%) answered “no”, and 15 (12%) answered “don’t know”.

**Figure 4.**
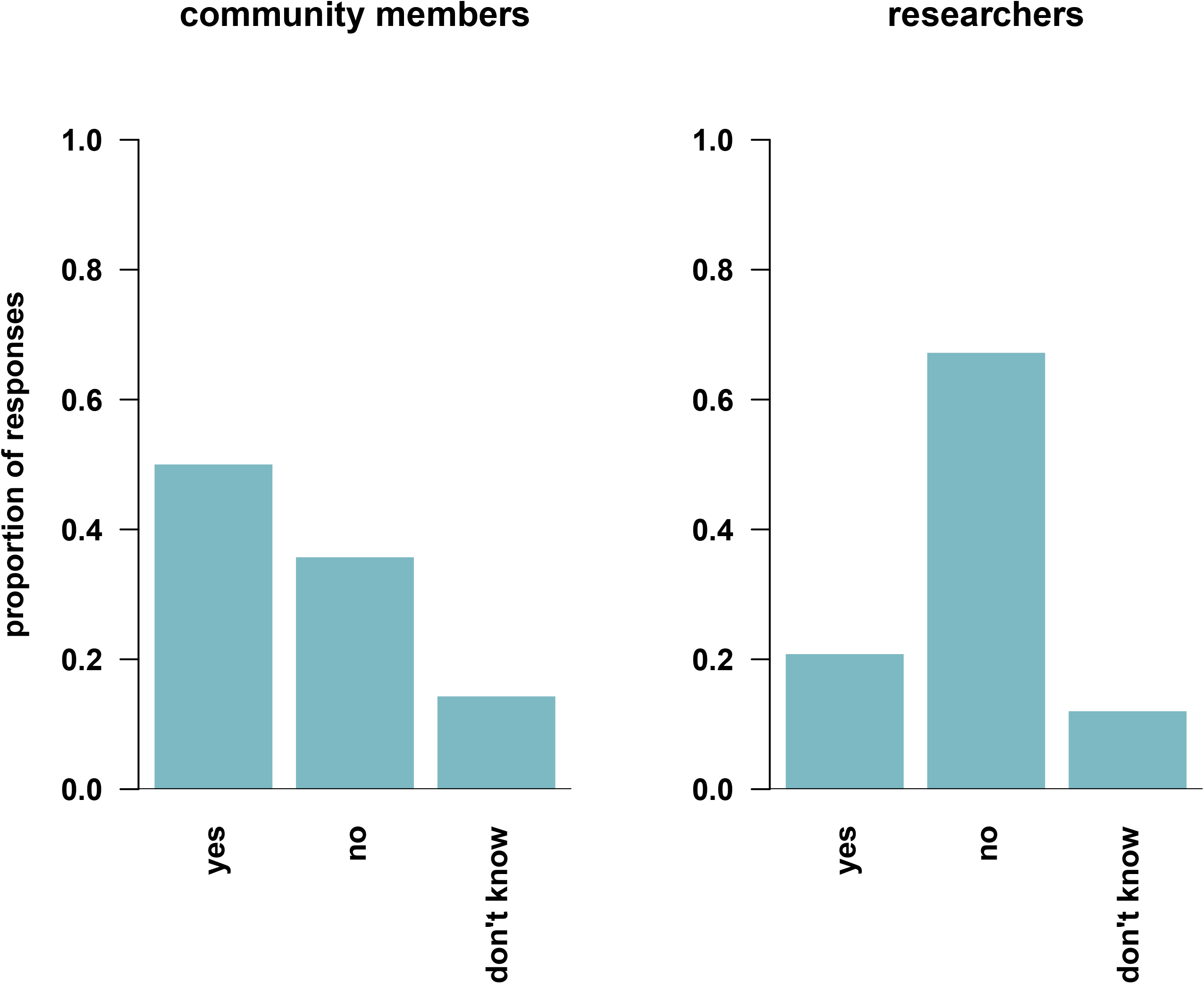
More Information. Survey participants were asked the structured question “Do you feel that you need more information about iPSCs before you can make an informed decision” with the following options: “yes”, “no”, “don’t know”, “do not want to answer”.

In addition to this multiple-choice question, all respondents were offered a free text field to share additional feedback to the following prompt: “Please share any additional information you would like to know about iPSCs.” Additional community comments focused on interest in more general information about iPSCs, more information about the types of iPSCs that will be shared, how they could be used in research, and the types of information that could be obtained from iPSCs, (see **Supplemental File C** for more detail). The majority of additional researcher feedback focused on interest in more technical and methodological information about potential NHGRI Repository iPSCs (see **Supplemental File C** for more detail).

## Discussion

Here we present results from an online survey regarding community and research perspectives on the potential creation of HapMap and 1000 Genomes Project iPSCs. The majority of community (86%) and research (86%) respondents agree that the relatively new technology now available to create iPSCs from HapMap and 1000 Genomes Project biospecimens holds potential benefits, including benefits involving differentiation into a variety of cell types and use in studying cell-specific functional implications of a wide range of well-characterized genetic and genomic variation.

Despite these potential iPSCs benefits, non-trivial concerns (36% community respondents and 16% researcher respondents) and doubts (21% community respondents and 11% researcher respondents answered “don’t know”) about HapMap and 1000 Genomes Project iPSC creation were shared (**Figure 3**). Of the five community respondents that reported concerns about the creation of HapMap and 1000 Genomes Project iPSCs, two respondents added additional free text feedback regarding concerns about the adequacy of the HapMap and 1000 Genomes project consent for iPSC creation. Similarly, of the 20 researcher respondents that reported concerns, nine shared additional free text feedback regarding concerns about the adequacy of the HapMap and 1000 Genomes project consent for iPSC creation: one of these respondents encouraged the use of more recently consented collections with explicit consent for iPSC creation; one respondent argued that the aim of the resource was focused on genetic data rather than phenotype data and iPSCs offer opportunities to collect cellular and developmental phenotype data that surpass the original consent; three respondents expressed concerns involving potential misuse of HapMap and 1000 Genomes Project iPSCs; one respondent argued that the lack of consent and potential for exploitation is even more concerning in situations that involve indigenous or marginalized communities; and one respondent expressed concerns about iPSC creation involving adequate protection for donors, their descendants and their communities.

Several research respondents shared free text feedback involving suggested limitations to sample usage. These include limitations to cloning, gamete production and usage, and use in transplantation or other clinical settings. We note here that the current NHGRI Repository governance forbids all of these uses for the current collection, which includes seven iPSCs that were created from newer donations from participants that explicitly consented to iPSC creation ^23^. Researchers (and their respective organization) requesting NHGRI Repository samples must formally agree to all of these restrictions in their agreement as well as to the additional following conditions of sample usage: not to attempt to identify or contact any donor or donor relative, not to use samples in human experimentation, and not to use samples in any therapies.

There are several limitations to our study. While we were not able to calculate a response rate for community members, we acknowledge that the community member feedback is incomplete, and we assume that our survey results do not represent the thoughts and beliefs regarding iPSCs of all of the donor community members. Contributing factors may include community member engagement attrition over time, challenges using Google Translate, and challenges accessing the online survey. In addition, given the age of the NHGRI Repository resource, which began distribution to the research community in 2006, we assume that we did not have active emails for all of the researchers that have received biospecimens over the past 19 years, and this may have contributed to the modest researcher response rate.

## Future Directions

Although both community and researcher perspectives were surveyed, the primary aim of our study was to invite feedback from the communities that have donated samples to the NHGRI Repository. The concerns shared by community respondents, which are echoed in the researcher feedback, preclude the creation of HapMap and 1000 Genomes Project iPSCs at this time. Given the potential benefits of incorporating iPSCs into the NHGRI Repository collection, we will continue to grow the existing iPSC collection ^23^ using new donor submissions through the Human Pangenome Reference Consortium (HPRC) ^24^ that explicitly consented to the creation of iPSCs. Similar to the 1000 Genomes Project, these new NHGRI Repository biospecimens will be extensively characterized through the HPRC with publicly available whole genome sequencing data generated with long-read and ultra-long-read sequencing technologies ^24^. In parallel, given the limitations of the online approach employed in the current study, we will continue to explore ways to connect with HapMap and 1000 Genomes Project community donors ^12^ that include the potential for in-person meetings to share educational materials and discuss views and understanding around the possible creation of iPSCs from their community biospecimens.

## Supporting information

Supplementary_FileA

Supplementary_FileB

Supplementary_FileC

## Acknowledgements

We would like to thank all of the communities and community members that so generously donated their samples and data for the creation of the NHGRI Repository. We would additionally like to thank all of the community liaisons, community members and researchers that took the time to share feedback through our survey. Finally, we would like to thank Alexander Arguello and Nicole Lockhart for their encouragement, feedback and support of this project.

