## Supplementary_FileA for "Community member, community liaison and researcher perspectives on the creation of HapMap and 1000 Genomes Project iPSCs"

### NHGRI Community iPSC Survey

**Welcome and thank you for taking this survey!**

We expect the survey questions to take between five and ten minutes to answer.

**Background**

The NHGRI Repository is a biobank that stores and shares DNA samples, cells and related information from people who took part in the International HapMap and 1000 Genomes Projects. People gave permission (consent) for their samples to be shared for many different types of research. The Repository does not share any personal or medical information to protect the privacy of the people that donated the samples.

Only researchers who meet specific requirements can access these samples. Researchers must submit a plan for their research study. If the research follows the consent rules, the researcher and their organization must sign an agreement to use the samples only for that study.

There is a new technology called induced pluripotent stem cell (iPSC) reprogramming. It can now be used on the samples donated to the HapMap and 1000 Genomes Projects. This means that the cells from the donated samples can be turned into different cell types, such as brain cells, liver cells, and heart cells. This gives researchers the opportunity to study different kinds of cells. For example, if someone wants to study how a specific heart disease medication works, it could be helpful to test the medication on heart cells made from iPSCs.

The iPSC technology did not exist when the HapMap and 1000 Genomes projects began, so participants were not told that their samples could be used this particular way. The original consent did allow researchers to use cells for many different types of research and for an unlimited source of DNA. However, creating different kinds of cells from their samples might not have been what participants expected.

We are doing this survey as part of a research study to learn what you think about using iPSC technology on HapMap and 1000 Genomes Projects samples. If you are willing to take part in this research study, please share your thoughts in this survey. Please with any questions.

**Do you identify as any of the following (please check all that apply): ***

- ☐ You have worked with one or more of the HapMap or 1000 Genomes Project communities to submit samples and data to the NHGRI Repository
- ☐ You are a member of a community that has participated in the HapMap or 1000 Genomes Projects
- ☐ You have used biospecimens from the NHGRI Repository for your research
- ☐ You have used genetic data associated with the NHGRI Repository for your research
- ☐ I am not sure
- ☐ Other

**Open text field answer:**

**Please tell us which population(s) you worked with. ***

You can select multiple options.

- [ ] African Ancestry in Southwest USA
- [ ] African Caribbean in Barbados
- [ ] Bengali in Bangladesh
- [ ] British from England and Scotland
- [ ] Chinese Dai in Xishuangbanna
- [ ] Chinese in Metropolitan Denver, CO, USA
- [ ] Colombian in Medellín, Colombia
- [ ] Esan in Nigeria
- [ ] Finnish in Finland
- [ ] Gambian in Western Division - Mandinka
- [ ] Gujarati Indians in Houston, TX, USA
- [ ] Han Chinese in Beijing, China
- [ ] Han Chinese South, China
- [ ] Iberian Populations in Spain
- [ ] Indian Telugu in the UK
- [ ] Japanese in Tokyo, Japan
- [ ] Kinh in Ho Chi Minh City, Vietnam
- [ ] Luhya in Webuye, Kenya
- [ ] Maasai in Kinyawa, Kenya
- [ ] Mende in Sierra Leone

- [ ] Mexican Ancestry in Los Angeles, CA, USA
- [ ] Peruvian in Lima, Peru
- [ ] Puerto Rican in Puerto Rico
- [ ] Punjabi in Lahore, Pakistan
- [ ] Sri Lankan Tamil in the UK
- [ ] Toscani in Italia
- [ ] Yoruba in Ibadan, Nigeria
- [ ] Do not want to answer

**Please tell us which community you are part of. ***

- ○ African Ancestry in Southwest USA
- ○ African Caribbean in Barbados
- ○ Bengali in Bangladesh
- ○ British from England and Scotland
- ○ Chinese Dai in Xishuangbanna
- ○ Chinese in Metropolitan Denver, CO, USA
- ○ Colombian in Medellín, Colombia
- ○ Esan in Nigeria
- ○ Finnish in Finland
- ○ Gambian in Western Division - Mandinka

- Gujarati Indians in Houston, TX, USA
- Han Chinese in Beijing, China
- Han Chinese South, China
- Iberian Populations in Spain
- Indian Telugu in the UK
- Japanese in Tokyo, Japan
- Kinh in Ho Chi Minh City, Vietnam
- Luhya in Webuye, Kenya
- Maasai in Kinyawa, Kenya
- Mende in Sierra Leone
- Mexican Ancestry in Los Angeles, CA, USA
- Peruvian in Lima, Peru
- Puerto Rican in Puerto Rico
- Punjabi in Lahore, Pakistan
- Sri Lankan Tamil in the UK
- Toscani in Italia
- Yoruba in Ibadan, Nigeria
- Do not want to answer

**Do you support the creation of iPSCs from your community's samples for research? ***

- Yes
- No
- Don’t know
- Do not want to answer

**Please share anything more about how you feel about the creation of iPSCs from your community’s samples.**

**Have you received any reports describing the ways in which your community’s samples are being used in research? ***

- Yes
- No
- Don’t know
- Do not want to answer

**Do you have any concerns about the way in which your community samples/data are being shared for research? ***

- Yes
- No
- Don't know
- Do not want to answer

**Please share anything more about the way in which your community samples/data are being shared for research.**

**Have you yourself ever contributed biological samples to the HapMap or 1000 Genomes Projects? ***

- Yes
- No
- Don't know
- Do not want to answer

**Do you support the creation of iPSCs from your samples for research? ***

- Yes
- No
- Don't know
- Do not want to answer

**Please share anything more about how you feel about the creation of iPSCs from your samples for research.**

**Do you support the creation of iPSCs from HapMap and 1000 Genomes Project samples for research purposes? ***

- Yes
- No
- Don't know
- Do not want to answer

**Please let us know why you do or do not support the creation of iPSCs from HapMap and 1000 Genomes Project samples for research.**

**Do you feel that there are potential benefits to adding iPSCs to the collection? ***

- Yes
- No
- Don't know
- Do not want to answer

**Please share anything more about potential benefits to adding iPSCs into the collection.**

**Do you have any concerns about adding iPSCs into the collection? ***

- Yes
- No
- Don't know
- Do not want to answer

**Please share anything more about concerns with adding iPSCs into the collection.**

**Do you feel that you need more information about iPSCs before you can make an informed decision? ***

- Yes

○ No

○ Don't know

○ Do not want to answer

**Please share any additional information you would like to know about iPSCs.**

**Please share any additional feedback you would like to include.**
