## Supplementary_FileC for "Community member, community liaison and researcher perspectives on the creation of HapMap and 1000 Genomes Project iPSCs"

**Free Text Survey Feedback**

Below is included the free text feedback we received from respondents. Each point is an exact quote that has not been altered in any way. Due to the relatively large volume of researcher free text feedback we have organized each item into sub-categories within each survey prompt.

Community Survey free text feedback to the following prompt: "Please let us know why you do or do not support the creation of iPSCs from HapMap and 1000 Genomes Project samples for research."

- iPSC holds tremendous promise for discovery and therapeutics and I do not believe that with proper safeguards, the donors and their communities would be harmed. That is why I support the creation of iPSC from the Yoruba samples.
- Potential of the repository to increase science knowledge.
- I'm not sure about supporting this because participants were not told that their samples could be used this particular way.
- A key finding of the community consultation with Indians in Houston in the early 2000s was that the community is generally open to the prospects of scientific discovery and research. Although iPSC creation was not envisioned at the time, science as such was regarded as a good and almost universally supported. I lend this my own personal support as an extension of and revalidation of that earlier finding. I do not believe that these fundamental attitudes have changed much in the years between when our community consultations happened and the present.
- It could be very interesting for research

Researcher Survey free text feedback to the following prompt: "Please let us know why you do or do not support the creation of iPSCs from HapMap and 1000 Genomes Project samples for research."

- Supportive feedback
    - This would be a really valuable resource of iPSCs for which we have many other datasets already (e.g. deep WGS, RNA-seq for many samples) plus representations of multiple ancestry groups that would be very helpful for genotype-phenotype mapping efforts in iPSCs.
    - Expands the range of quality tools in human in context of reduced access to (and confidence in) animal models.
    - Would be useful for comparing different cell types from the same individuals. Could facilitate cellular GWAS, which we've done previously only with LCLs at this point. We're going with organoids in our own work right now, but iPSCs would allow a broader range of cell types,.
    - Because they are well characterized genetically, any phenotypic results obtained could potentially be traced back to the related genes/proteins . This information might not be available from other sources. Understanding the phenotypes or failure to produce them could be essentail to understand whether to use IPSC tech on humans, organoids, or other therapeutic applications.
    - genetic diverse iPSc with sequence information is very valuable.  There is already so much information about these samples so iPCs will allow the community to build on those knowledge.
    - Combining the genetic background we already have with phenotypic and molecular profiling data would be invaluable.

- It is a good idea to have iPSCs from characterized samples. This is less critical as iPSC generatoin becomes easier from samples that are clinically relevant. The utiolity of specific cells will depend on the quality of the WGS an the metadata
- It would provide an additional set of iPSCs with matched genomic sequences to the research community.
- Having worked with iPSCs in the past, I am aware of their versatility and utility in human genomics research.
- This would be a great resource for studying the effects of genetic variation on complex disorders that can be modeled in hIPSC derived tissues
- A valuable tool for researchers with tissue-specific needs
- Correlate function with huge amount of genetic data available
- This valuable resource can provide tailored iPSC lines to a broad spectrum of researchers that may not have the resources to generate the iPSC lines themselves.
- Multi-omics data can be generated for these samples as controls.
- Highly valuable resource, especially given the open consent for genomic sharing from these individuals. There is an opportunity to provide a critical need to the community here and solve a true pain point.
- It will generate new materials for research by organ type
- Could be useful for relating genetic variation to functions in specific tissues by differentiating the iPSC into specific lineages
- there is a lot of genetic data known about these cells, so generating tissue specific models would leverage all that data for multiple uses.
- Having hiPSC resources where there is complete or near-complete haplotype information will be invaluable for research aiming to make genotype-phenotype infrerences
- It would be very helpful to have diverse iPSCs that could be transformed into different cell types for functional genomics validation of functional variation.
- Additional lines to make mutations against an isogenic background woul be very valuable.
- The genetic resource of hapmap/1kg is outstanding, but LCLs are the obvious limitation to studying genetic effects. IPSCs would expand that range exponentially!
- Moving scientific research forward
- We published a study on variants Affecting mRNA splicing in the samples. It would be worthwhile to determine whether the allele specific isoforms we observed are tissue specific or agnostic.
- IPSC from reference material is rare and we know the Hapmap and 1000 Genome well.
- could use them to phenotype and perform cellular GWAS of human traits
- An iPS cell line with precise genome data is precious and inevitable resources for biomedical research and development.
- potential for therapeutic benefits
- Invaluable tissue specific information for new technologies that will allow for standardized materials
- Most of all I would appreciate to use iPSCs from HapMap pedigree 1463, because the entire family has been sequenced T2T recently. Thus, these are the best characterized human genomes on this planet.
- The iPSC cells would be a valuable resourse for our research.

- We will have high-quality genomes of these individuals, with genetic differences that could affect the basic function or disease in different cell types. You can only determine if different differentiated cell types are affected by converting the LCLs to iPSCs.
- For the rare diseases they provide the only source of QC or validation material/reagent
- iPSC from different human ancestries are very valuable, connection to health data would be valuable as well
- Different genotypes are found in the different populations
- It will hopefully generate consistent cell lines to help standardize research results
- these people are incredibly well genetically characterised and being able to cherry pick a particular genotype for iPSC without having to recruit your own participants would be great
- The diverse genomic information offered by the projects will contribute to advances in science significantly
- iPSCs could be used for more studies that require specific cell types.
- Do support. Provides a readily available resource for boosting many scientific otojects
- The genetic diversity of the HapMap and 1000 genomes samples would allow for the selection of optimal genomic backgrounds and optimal controls for functional analysis of iPSC derived cell types and organoids.
- The genetic variation that is inherent to the HapMap and 1000 Genomics Project is an invaluable resource for all researchers in Genetics and Genomics.
- seems like generation of this data will provide a generally useful and important data set for the future
- It will be very useful to know the exact genetic background of differentiated iPSCs for disease research.
- it is always valuable to have different material to use in research.
- It will facilitate our progress in the field of biology.
- Having iPSC of individuals with specific genetic variants identified in assemblies of high-quality and complete, is an extremely powerful resource for precision medicine and basic scicence.
- I think the HuBAMP lines will be quite valuable for researchers. They will derive organic models from these.
- to study genome diversity and phenotype
- Much of the work that we do relate to brain phenotypes making LCLs not amenable to molecular results. Being able to differentiate iPSCs to neural cells (and others) would be valuable.
- The samples are useful for confirming gene editing activity at variant sites that are present in the population at relatively low frequency. However, editing may not occur in lymphoblastoid cells carrying such variants due to chromatin accessibility issues in that particular cell type. The availability of iPSCs would enable the cells to be reprogrammed to the relevant cell type to enable editing of variant cell types to be investigated in a more biologically/therapeutically relevant manner.
- It would be very useful for testing genetic hypotheses in the lab

- Scientific concerns / limitations
  - Don't know what the end product would be and how useful they would be for research. They would need to be characterized.
  - I don't think there would be great value, as there are now many better resources than HapMap/1KG for such activities that have linked phenotypic and clinical data and have been fully sequenced

- Although these can be different types of cells, they are still generally individual cells or small collections of cells and thus are not that different from ordering a lymphoblastoid cell line or a fibroblast cell line.
- They must be from primary fibroblast punches. Do not reprogram the LCLs. The lab that performs the reprogramming is important too.
- These lines can potentially serve as controls for studies of iPSC lines from study participants with candidate mutations. For the NHGRI iPSC lines to be useful, it will be critical to know that the donors are true controls; that is, with no severe genetic conditions.
- genomic QC will be vital
- I support the creation of iPSCs from these samples but there is a need for a discussion as to the protocols used and how to avoid confounders in cross-population comparisons.

- Ethical concerns
  - have no objection for the creation, but not sure ethically. so far research concerns, it should be done.
  - I have reservations about this being done in the absence of explicit consent from the original participants, particularly for anyone from indigenous communities.
  - Despite their potential value, I have concerns about ethical overreach.
  - Participants were not asked for that use
  - If we create appropriate protection measures for the communities and descendants of the people that donated samples for the HapMap and 1K Genomes project I would consider this to be a great tool to help us understand functionally health and disease
  - Yes, it would be useful, but I am uncertain if the consent supports that activity
  - More recently consented cell collections we have worked with are explicitly consented for iPSC creation or use, so I am hesitant to say whether I think this would be an acceptable use within the existing consents. If it were decided that this is acceptable, I would imagine we would take advantage of the resource, but I'm not going to say I think that's reason enough to do it. I think a related question would have to be whether the use of iPSCs would include the ability to edit the lines, another technology which was not anticipated in the original consents.
- Other
  - I am not doing the research on cell line, but I think it would benefit for others
  - I don't have enough information to form an opinion
  - This use would fall within what a reasonable submittee might expect their samples to be used for so in most cases would be welcomed and rarely, if ever, opposed. The burden of reconnecting for this specific use seems far too high for any benefit beyond the social aspect of dotting every possible I.
  - My sense is that a broad consent to use patient cells would include a manipulation of these cells that enables analysis of a variety of cell types.
  - All of my research for the past 50 years has been genomic-based
  - Support because it will make a central deidentified and approved repository with appropriate consenta
  - The donors have not been collected for commercial purposes, which is limiting the scope of the project. Otherwise, it's a great resource for R&D!
  - We usually do not utilize iPSCs for any projects.
  - This questionnaire does not include statements of global agreement, consensus and collaboration policies with other institutions, such as the Center for iPS Cell Research and Application (cira), Kyoto University which has been contributing to the development

of iPS cells. HapMap and 1000 Genomes Project samples were collected worldwide, which means that global discussion and consensus are needed to develop and meet worldwide expectations.

- I have insufficient knowledge on the cost/benefit ratio
- It is not necessary to make iPSCs from all samples but a subset would be valuable
- My research focuses on genetic screening, not in the field of stem cell technology.
- No core interests in iPSC-based research
- My work does not involve iPSCs but I could see how they could be useful.
- Different cell types can respond very differently to treatment. I think that participants who agreed to let their blood cells be used in various types of experiments would agree as well to let their samples develop different cell types.
- not sure it will apply to my area of research in cancer
- It might be more cost-effective to freeze cells from subjects that could be turned into iPSC if needed rather than make them all at the start
- I don't see how this would be different than people using the LCLs
- I do not have technology to do so.
- No a need in my research at this time.
- unsure what the use case is vs other cell lines that are available or can be acquired. the ease of generating DNA data makes other issues, such as line quality etc more important
- NHGRI has already transformed cells in a sense by creating LCL's. Inducing pluripotency to create different kinds of cells is in the same general class of transformation that HGDP donors consented to. Moreover, they consented to broad use of their DNA, and understanding how their DNA shapes the regulatory environment of their cells of different types may be construed as being within the original consent.
- It would be a great resource for other scientists' work (I just happen not to need it for mine)
- An easy accessible, source of healty iPSCs of human populations that are genetically characterized is very helpful; however whether this is done best with the HapMap samples is not entirely clear to me
- The subjects donated their samples for research and iPSCs are consistent with that original consent.
- No opinion.
- Seems like a large expense that would be utilized by very few. Centrally supported projects should benefit a very high percentage of investigators. This project does not meet this standard.

Community Survey free text feedback to the following prompt: “Please share anything more about potential benefits to adding iPSCs into the collection.”

- Adding iPSCs into collection will be important for addressing challenges with available diverse iPSC resources that biomedical research needs to make research outcomes more generalizable.
- Enables in vivo studies.
- New applications or additional studies can be performed providing new solutions for diseases
- This new technology offers the opportunity to improve research and it is a challenge.
- Possibility of different cell types, with genetic information
- IPSCs from the 1000 genomes projects were fully genetically characterized at no additional cost to the investigators.

Researcher Survey free text feedback to the following prompt: “Please share anything more about potential benefits to adding iPSCs into the collection.”

- Scientific benefit
    - Matching genotypes to phenotypes
    - With the extensive genetic data available the success and function of the iPSCs could be far better understood.
    - A useful service for the research community at large to use as standards.
    - Unlike LCLs, these are not transformed and should not have acquired genetic abnormality during culture
    - I think the generation of iPSC's for rare diagnoses may have a higher impact than 1000 Genomes samples. However, having the 1000 Genomes sample iPSCs would serve as an excellent control for rare disease lines.
    - Some genes can be differentially expressed in different tissues - this may be a way to get a better understanding of differential splicing and differential expression
    - In my case,pharmacogenomics, this would greatly broaden the array of drugs and organs that we could study.
    - A one-stop shop for custom iPSC lines with valuable metadata availability.
    - Valuable public resources
    - monitor changes in that occur from the induction that will identify regions of interest for further study
    - It should facilitate discovery of new therapeutic targets.
    - Could be useful for relating genetic variation to functions in specific tissues by differentiating the iPSC into specific lineages
    - For the global genomic variation of the HapMap and 1000 Genomes Projects, there are two potentials to contribute to plantation medicine for different types of HLAs and to life sciences aiming to reveal global variation in development and gene expression.
    - Carrying out fundamental genetics of human cell biology and development using iPSC-derived cells and organoids, with open data release.
    - This will enable research to develop differentiated cell lines to study natural human variation and disease-causing mutations in cells that are relevant models of specific human tissues. The proposed iPSCs will have many applications in basic science and medicine.
    - Genetic variants that impact iPSC properties
    - Most of all, differentiation of these cells into cell types of interest would provide a standardized control for transcription analyses.
    - Can study important biological questions in different cells types in culture.

- Being able to request lines with a specific variant of interest
- These lines can potentially serve as controls for studies of iPSC lines from study participants with candidate mutations. For the NHGRI iPSC lines to be useful, it will be critical to know that the donors are true controls; that is, with no severe genetic conditions.
- It would promote basic research of iPSC.
- gain functional understanding of the effect of variants associated with different diseases in different populations around the world
- There are plenty of position papers in the literature (e.g. PMID: 37708421) affirming the value of iPSC.
- Providing iPSCs from the collection would enable some researchers access to a valuable resource they might not have the opportunity otherwise.
- These iPSCs will have matched high quality assemblies
- The addition of iPSCs will allow researchers to test hypotheses related to the impact of genetic variation on how different cell types express specific RNAs and proteins.
- don't even know where to start, but cell types matter a lot when one wants to understand genotype-phenotype relationships and iPSCs is the most powerful way to get different cell types from one renewable resource read
- I think the ability to leverage the vast amount of knowledge we have on the genetics of these individuals and the ability to pick specific variants on a natural background across diverse populations would allow many genotype-to-function studies that would require more work without a well-characterized resource like this.
- Support cell therapy and generative medicine
- iPSC can be differentiate in different cell types, and the variants of different individuals can affected those cell types differently.
- diverse iPSC sets (different genotypes, disease states, etc) can help basic and translational research
- add additional epigenetic information on top of genome information
- So a diversity of cell types could be created to understand haplotype/genetic effects in different cell contexts.

- Other
  - may be this will benefit, not sure about acceptance in the population.
  - Serach Criterial and the display should have more interactive and desriptions.
  - I don't have enough information to form an opinion
  - We can save a lot of money, labor, and time
  - adding iPSC with commercial intent for CRO and pharma companies to use as well.
  - it would save investigators a lot of time and money.
  - Do it!
  - It is beneficial to adding iPSCs into your portfolio.

Community Survey free text feedback to the following prompt: “Please share anything more about concerns with adding iPSCs into the collection.”

- A major concern is getting permission to create iPSCs from the samples if this was not in the initial consent used for collecting the samples.
- International guidelines, standards and registries should be used as reference
- Lack of inform consent from donors.

Researcher Survey free text feedback to the following prompt: “Please share anything more about concerns with adding iPSCs into the collection.”

- Consent related
    - You need ironclad consent to do this.
    - Pros and Con of having iPSCs, any risks of having it, consent of using the sample.
    - As above, concerned about extent of original consents, and perceptions of exploitation of tissues and data particularly for indigenous or marginalized communities
    - The consent for cell line creation was in order to provide a renewable source of DNA. The aim for both sample sets was explicitly to provide genetic but not phenotype data. Turning them into iPSCs would only be in order to collect cellular/developmental phenotypes. This transition to phenotype-based genetics goes fundamentally beyond the original consent concept. We should use cell lines explicitly consented for cellular phenotype research for these purposes.
    - The main issue is an ethical one--whether or not it was covered by the original consent form.
    - I would want to be sure the subjects were properly consented for iPSCs.
    - Consistent with consent?
    - There are potential uses for iPSC which the donors may not have approved if asked: cloning, gamete production (and use), and allogeneic transplant. I would suggest creating iPSC but limiting use to the least objectionable possibilities, i.e. creation and assay of cell types and organoids for research use only.
    - As long as the contributor has provided the appropriate consent, this is a fantastic resource.
    - I am concerned that the consent was not designed to support iPSC establishment and biobanking
    - This may go beyond the research for which these groups consented.
- Other ethical concerns
    - I am concerned about misinterpretation about social discrimination of research results using global samples. I am also concerned about the chaotic use of iPS technology, which has been carefully managed by the cira of Kyoto University.
    - Possible misuse or manipulation, more complexity in regulation, and public mistrust.
    - I am concerned about the protection to descendants of the donors of this projects and also to the communities.
    - Potential use in unethical research
- Scientific / methodological concerns
    - Need to have WGS with the cells or at least the donors
    - This activity will do more harm than good to research if some of the lines are from people with damaging genetic lesions. It's important that these people be true controls.
    - will they be sequenced to check no major acquired mutations

    - Cell line research is replete with artifacts. Recognizing these limitations can be a challenge.
- Other
    - I don't see iPSCs any differently that LCLs
    - I don't have enough information to form an opinion
    - Not if carefully regulated and vetted for quality research programs.
    - I am sure it this will be considered, but uniform reprogramming protocols must be used. Variability in LCL creation has caused issues with current 1KG samples and current iPSC collections also suffer from these artifacts.
    - They must be from primary fibroblast punches. Do not reprogram the LCLs. The lab that performs the reprogramming is important too.
    - you need to be clear about exactly what criteria can be used to cherry-pick samples
    - Expense
    - Only cost
    - People won't stop using iPSCs anyway. Therefore having iPSCs that can be used by many labs will establish a common platform
    - the samples are anonymous and reconsenting is not possible. the broad original consent can be considered to apply

Community Survey free text feedback to the following prompt: “Please share any additional information you would like to know about iPSCs.”

- Types of iPSCs that will be shared and their use.
- I will like to have more information about what it means to "add iPSC s to the collection". The meaning of this is not clear to a layman and that is why I answered don't know in these sections.
- Patientes should be previously informed about this technology
- What type of experiments could be carried out with iPSCs and what information could be obtained with this technology.
- I do think that an information sheet would be good to circulate. the survey intro as such provides only scant information -- many may be compelled to search online before responding.

Researcher Survey free text feedback to the following prompt: “Please share any additional information you would like to know about iPSCs.”

- sample characteristics
- Is creation of this resource consistent with how samples were consented?
- Pros and Con of having iPSCs, any risks of having it, consent of using the sample.
- I don't have any information currently
- We are a stem cell company. No concerns here! Only praises!
- Ethical and legal agreement and global concensus, as I mentioned earlier.
- None.
- N/A. My work does not involve iPSCs.
- I have no problems in allowing to create iPSCs to a new collection of samples were consent have been given for that but not in the old sample sets as they are.
- Is there health history data available on this lines
- characterization and quality control of iPSC.
- More information about the legal backbone that will facilitate the research and protection of human subjects will be essential for the development of this tool
- Regulations on access to iPSCs
- HGDP iPSC will be derived from LCLs.  I would want to know if they will therefore be preserving artifacts of the LCL transformation.
- Review of the consent.
- genomic analysis
- iPSCs are only useful when they work well which means they need to be characterized substantially; not sure whether the HapMap cell lines are the best source for making iPScs
- which protocols will be used for reprogramming; how to ensure that the communities who donated their samples will be protected from unlimited use of the iPSCs given that we don't know the full range of possible uses of these lines.
- Process of reprogramming that lead to the iPSCs
- PIGI consent, patient information (age, sex, disease type or healthy), methods used to collect patient cells and generate iPSCs, karyotype
- No opinion.
